# Attention level assessment by means of HRV data extracted from fNIRS signals

**DOI:** 10.64898/2026.02.02.703265

**Authors:** Mahya Sadat Aramoon, Seyed Kamaledin Setarehdan

## Abstract

Sustained attention is an important requirement for high performance in all cognitive processes. Quantifying the level of sustained attention to prevent attention lapses is therefore necessary for effective human-machine interfacing. Furthermore, sustained attention evaluation can help diagnose and treat attention deficit hyperactivity disorders. Attention level can be assessed by brain and heart signals. This study employed functional near infrared spectroscopy (fNIRS) and the heart rate variability (HRV) information extracted from the fNIRS signals to differentiate the rest and three levels of sustained attention states. Sustained attention states are induced by three modified versions of continuous performance tests (CPT). Eight subjects engaged in three sessions of attention tests. fNIRS brain signals were recorded from the right prefrontal and dorsolateral prefrontal cortex. HRV information was then extracted by processing the fNIRS signals. For attention classification, support vector machine (SVM), linear discriminant analysis (LDA), and random forest (RF) algorithms with mutual information based feature selection were applied on the fNIRS and HRV data both separately and together. In the classification of the three levels of attention using fNIRS and HRV data, the LDA classifier showed the best performance accuracy of (80.9 ± 1.5%) and (56.2 ± 1.0%), respectively. For two-class classification between the rest and the attention states (all together), the accuracies of (98.9 ± 0.3%), (95.6 ± 1.2%), and (99.5 ± 0.2%) were obtained using the RF classifier on the fNIRS, HRV, and combined data, respectively. These results demonstrate the effectiveness of the HRV data for classifying sustained attention states. Moreover, using the combined fNIRS and HRV data provides better classification accuracy.

## Introduction

Sustained attention is the ability to keep attention on the stimulus for a long time, playing an important role in memory^1,2^ and learning^3^ mechanisms. Sustained attention assessment and preventing attention decrement have a great significance in human life activity, such as driving, Air traffic control, navigation, and other human-machine interfaces^4^. Furthermore, evaluating sustained attention can help diagnose and treat attention disorders such as attention deficit hyperactivity disorders (ADHD), who cannot keep their attention continuously on one task and are distracted easily by irrelevant information^5^. So sustained attention evaluation has a benefit for both patients and healthy people’s lives.

Behavioral measurements estimate the sustained attention levels and vigilance during cognitive tests such as continuous performance test (CPT)^6^, psychomotor vigilance task (PVT)^4^, and Sustained Attention to Response Task (SART)^7^. CPT is the most common test widely used in neuropsychological research like brain damage, schizophrenia, ADHD, and healthy people for investigation neural basis of attention^6^. CPTs have different versions, which are causing various levels of attention. They are divided into three main categories including CPT-X, CPT-AX, and CPT-IP^8,9^. In all version of CPTs, the stream of the stimuli are shown to participants, and they should respond to the targets as quickly as possible. During the test rate of true answers, false alarms and reaction time for each participant were recorded for evaluating the attention level.

Besides behavioral performance, physiological measurements can assess sustained attention. in past studies, electroencephalogram (EEG)^10^, functional magnetic resonance imaging (fMRI)^2^, and functional near infrared spectroscopy (fNIRS)^11–13^ have been used for measuring brain signals to investigate the neural basis of attention. fNIRS is a new optical device for measuring oxygenated hemoglobin (HbO) and deoxygenated hemoglobin (Hb) changes in the brain cortex^14^. Since sustained attention is originated from the prefrontal cortex (PFC) and dorsolateral prefrontal cortex (DLPFC), fNIRS is an appropriate device for monitoring the hemodynamic response during attention tests. In Recent studies, fNIRS signals have been used to discriminate attention levels^4,15,16^. Furthermore, researches show the right PFC and DLPFC are more activated than the left one during sustained attention^11,13^.

Heart signals also evaluate sustained attention. The autonomic nervous system regulates involuntary functions in the body, such as the heartbeat via the sympathetic nervous system (SNS) and parasympathetic nervous system (PNS). There is an integration between mental state and autonomic function represented by the “neurovisceral integration” (NVI) model. In this model central autonomic network (CAN) in the brain modulating heart rhythms by controlling sympathetic and parasympathetic neurons. CAN includes several regions in the brain such as the medulla, amygdala, and prefrontal cortex. In this network, the medulla integrates information of efferent nerves that come from the brain and afferent nerves that come from the heart. During upward and downward pathways in CAN, the cognitive and attentional processes in the PFC and heartbeats can affect each other^17^. heart rate variability (HRV) has been used to represent this relationship. HRV is the variation of the beat-to-beat intervals, which are measured by the time between two consecutive R peaks (R-R intervals) of the electrocardiogram (ECG) signal. Past studies suggest that higher HRV is related to better performance in a cognitive task such as working memory, attention, and inhibitory control^18–20^. Also, the high-frequency and low-frequency power of HRV decreased during attention tests^21–23^. In addition, HRV has been used in biofeedback systems for improving attention. For example, in brain-damaged patients increasing the low-frequency power to high-frequency power ratio of HRV caused greater performance in the attention task^24,25^.

The fNIRS measures the hemodynamic response which is contaminated by physiological noises such as the heartbeat, respiration, and Mayer waves and nonphysiological noises such as motion artifacts and optode movements. Several studies have been done to extract heartbeats from fNIRS signals for heart rate (HR) analysis^26,27^. Recent one suggested the real-time mean weighted algorithm consisting of four peak detection algorithms including AMPD^28^, S1 and S5 Functions^29^, and M2D^30^ with the specific correction process effectively extracts HR^31^. Hence, by extracting HR extraction from fNIRS signals, two functional information sources from the brain and heart are provided simultaneously.

Although previous studies investigated sustained attention by fNIRS and HRV signals, none of them considered both of these signals simultaneously. HR derivation from the fNIRS signals allows having both the brain and heart responses with one fNIRS device during attention.

In most research, the sustained attention level has been classified into the low and high attention levels with fNIRS or HRV signals individually. Only a few studies have done more than two levels, but they could not reach acceptable accuracy. Hence, it is necessary to find the best discriminative features related to sustained attention in fNIRS and HRV signals and combine them to determine attention levels more accurately. In summary, the current study’s objective is to investigate and compare the potential of fNIRS and HRV data to classifying different levels of sustained attention and determine the optimal combination of features for classification. This study has a particular advantage over other studies because it used HRV extracted from fNIRS signals to obtain both brain and heart responses with one device. We designed three modified versions of CPT-X, CPT-AX, CPT-IP-to induce three sustained attention loads. For increasing the quality of fNIRS signals, some signal processing algorithms, including the wavelet filtering^32^, spline interpolation^33^, bandpass filtering, and principle component analysis (PCA)^34–36^ were utilized. Support vector machine (SVM)^37^, linear discriminant analysis (LDA)^38^, and random forest (RF)^39^ were used for classification. Due to the importance of selecting the most discriminative features, the mutual information based algorithm was applied. Since the hemodynamic and heart responses dependent on individual characteristics, similar to previous studies, classifications were done on each participant and averaged accuracy over participants was reported^4,40–46^. As it was expected, HRV features can classify sustained attention levels, which confirmed the connection between heart and brain, but fNIRS features have better classification performance. Moreover, combining HRV and fNIRS features caused more classification accuracy in all classifiers.

The rest of the paper is organized as follows. In section 2, the materials and methods are explained. In part 3, after presenting the results of behavioral measurements, attention classification results are demonstrated. Section 4 and 5 are describing the discussion and conclusion, respectively.

## Materials and Methods

### Participants

Eight healthy, right-handed, female students (aged 23.8± 0.97 years) from the University of Tehran participated in the experiment. Based on the self-report, they had normal vision and no history of psychological, neurological, and cardiovascular disorders. After describing the experimental procedure, written consent was obtained from all of the participants.

### Signal acquisition

The continuous wave fNIRS system developed at the University of Tehran^47,48^ with wavelengths of 730 nm and 850 nm was employed for measuring the brain hemodynamic response. Eight fNIRS channels, including two light sources and six detectors, were placed on the right PFC and DLPFC. Data were recorded at a sampling rate of 10 Hz, and the source-detector distance was set to 2.1 cm. The modified Beer-Lambert law was used to convert raw intensity signals to the HbO and Hb changes^14^. Figure 1 illustrates the configuration of sources, detectors, and channels.

**Figure 1.**
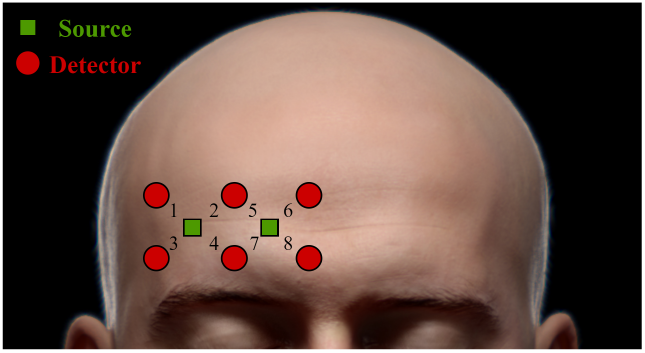
The placement of fNIRS sources, detectors and eight fNIRS channels on the right PFC and DLPFC.

### Continuous performance tests

For evaluating sustained attention, CPTs in three levels of difficulty named CPT-X, CPT-AX, and CPT-IP were designed by PsychoPy library^49^ in python with the following description.

- CPT-X: in this test, the participants were shown a random stream of different letters. They had to respond to the target letter “X”. 25% of stimuli were target trials. It is the easiest version of the CPT and causes less attention load.
- CPT-AX: Like CPT-X, a random sequence of different letters was shown, but participants had to respond to the letter “X” which goes immediately after the letter “A”. In this test, trials consist of “X” letter following an “A” letter (“AX” trials (targets), 25% pairs), non-X letters following the “A” letter (“AY” trials, 33% pairs), the “X” letter following non-A letters (BX trials, 28% pairs) and non-X letters following non-A letters (“BY” trials, 14% pairs).
- CPT-IP: this test is more complex than CPT-X and CPT-AX. In CPT-IP, in each trial, the stimulus was a string of three letters. Subjects were instructed to respond to the stimulus which was identical to the previous one. Two successive trails had three types. 25% of them were identical pairs (target). 45% were catch-pairs in which the order of letters in the string or one letter in the string was different between two trails. The remaining pairs were utterly different from each other.

Each CPT consisted of 90 trials in which the stimulus appeared in the middle of the screen in white color. The stimulus was presented 100 ms and followed by an interstimulus interval (ISI) of 1500 ms or 2000 ms, with each ISI appearing in 50% trials. Participants were informed to respond target by pressing the space bar as soon as the target appeared. Figure 2 depicts the schematic of the CPT-X, CPT-AX, and CPT-IP paradigm.

**Figure 2.**
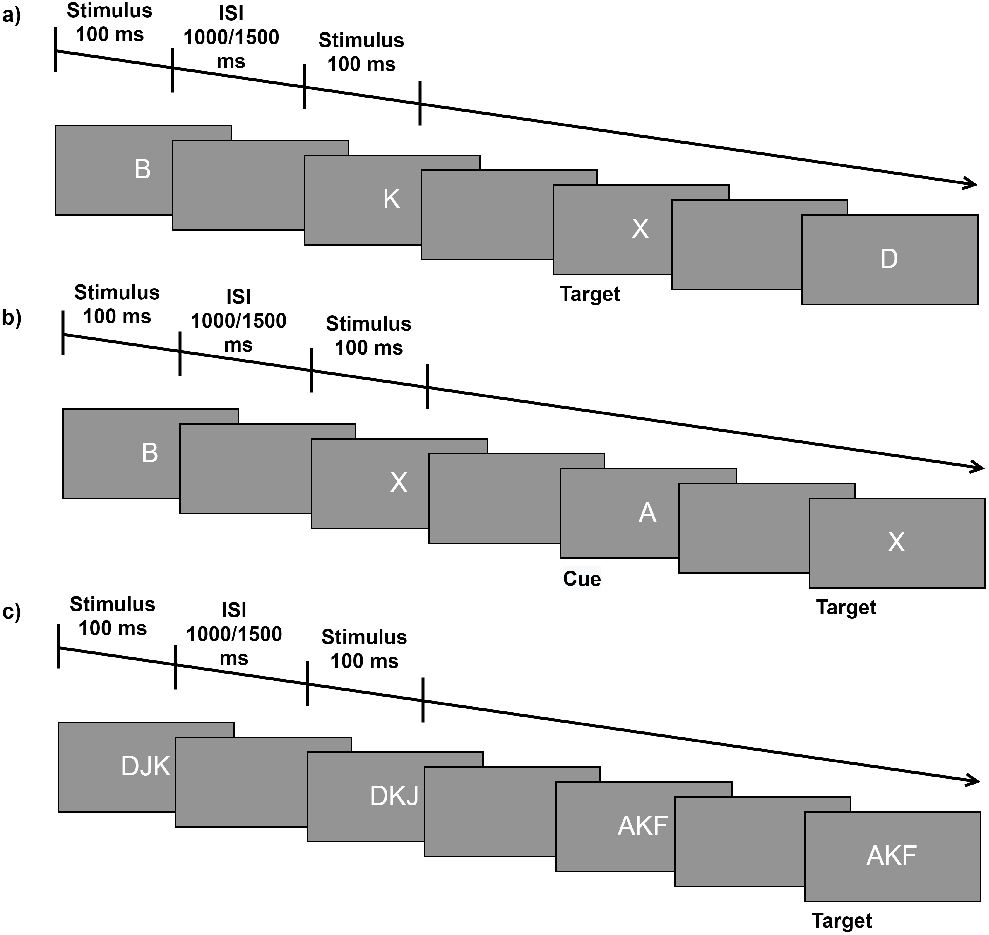
An example of the trial sequence of a) CPT-X b) CPT-AX c) CPT-IP. Each stimulus lasted 100 ms, and ISI was chosen randomly between 1000 and 2000 ms.

### Experimental paradigm

The experiment was done in a silent, ventilated room with appropriate lighting. The participants were asked to sit on a comfortable chair placed approximately 80 *cm* from a 19.3*34.4 *cm*^2^ monitor. They were also advised to be relaxed, silent and avoid head movements during the test.

Three test sessions were considered for evaluating three different sustained attention levels. In each session, participants performed one of the CPT versions (CPT-X, CPT-AX, and CPT-IP). one session lasted 705 s, and there were 900 s of rest sessions between two consecutive sessions. To avoid adaptation, each participant conducted three CPT sessions in random order.

At the beginning of each test session, participants received a complete description of the test then they performed test sessions according to the experimental paradigm shown in figure 3. One session included three blocks of test which were separated by 30 s rest blocks. In the first and last part of the session, there were 150 s and 120 s rest blocks, respectively. The preparation blocks were used to inform the participant that the test starts in few seconds.

**Figure 3.**
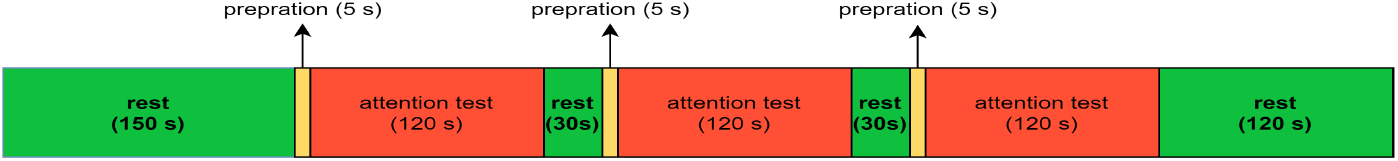
Schematic illustration of experimental paradigm in each session: The green blocks represent the rest periods; the red block represents the test period; orange blocks represent the preparation period for preparing the participant for the test.

The rate of true answers, false alarms, and the averaged reaction time during the test were recorded to evaluate participants’ performance. Furthermore, at the end of each session, participants were reported their attention score with a number between 1 to 10.

### Proposed method

Figure 4 illustrates four main sections of the proposed method which are preprocessing, HRV derivation, feature extraction, and attention classification. Due to poor signal quality, channels 3 and 6 of the fNIRS data were excluded from further analysis.

**Figure 4.**
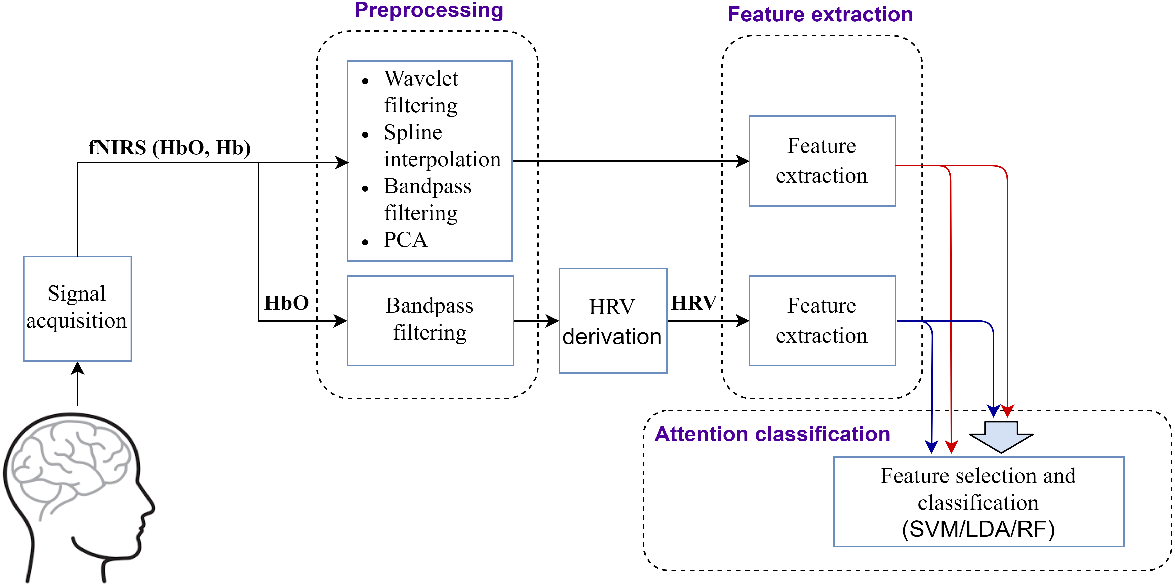
Block diagram of the proposed method for attention classification.

### Preprocessing

In order to extract hemodynamic response, three preprocessing steps were applied to reduce noise and artifacts from fNIRS signals of all subjects. In the first step, wavelet filtering described by Molavi and Dumont^32^ was used to remove abrupt changes in signals due to decoupling of optodes from the scalp during sharp motion artifacts^50^. The second step, moving standard deviation and spline interpolation proposed by scholkmann et al. was employed to eliminate baseline shift and slow motion artifacts^33^. In the third step, a zero-phase low-pass FIR filter at 0.004 Hz and a zerophase high-pass FIR filter at 0.15 Hz was applied to remove physiological noises and baseline drift^51^.

After applying three main preprocessing steps, two subjects had low periodic oscillations that appeared in all their channels. To remove this artifact, the PCA algorithm^34^ was applied on fNIRS signals. Since this artifact appeared in all channels, the first component of PCA represented the artifact variation. Hence, the artifact was eliminated by removing the first component and reconstructing signals. After preprocessing, HbO and Hb were averaged over all channels^11^ then the mean of the first rest block was subtracted from the whole signal. Averaging across all channels increases reproducibility of fNIRS measurements^51,52^. Figure 5 shows the grand averages of fNIRS signals and the standard error of the mean (SEM) during CPT, CPT-AX, and CPT-IP. In this study, the grand average is the average of blocks across all channels over all participants. According to figure 5, the amplitude of HbO changes in CPT-IP was more than CPT-AX, and the amplitude of HbO changes in CPT-AX was more than CPT, which is proving designed CPTs evoked a different level of sustained attention.

**Figure 5.**
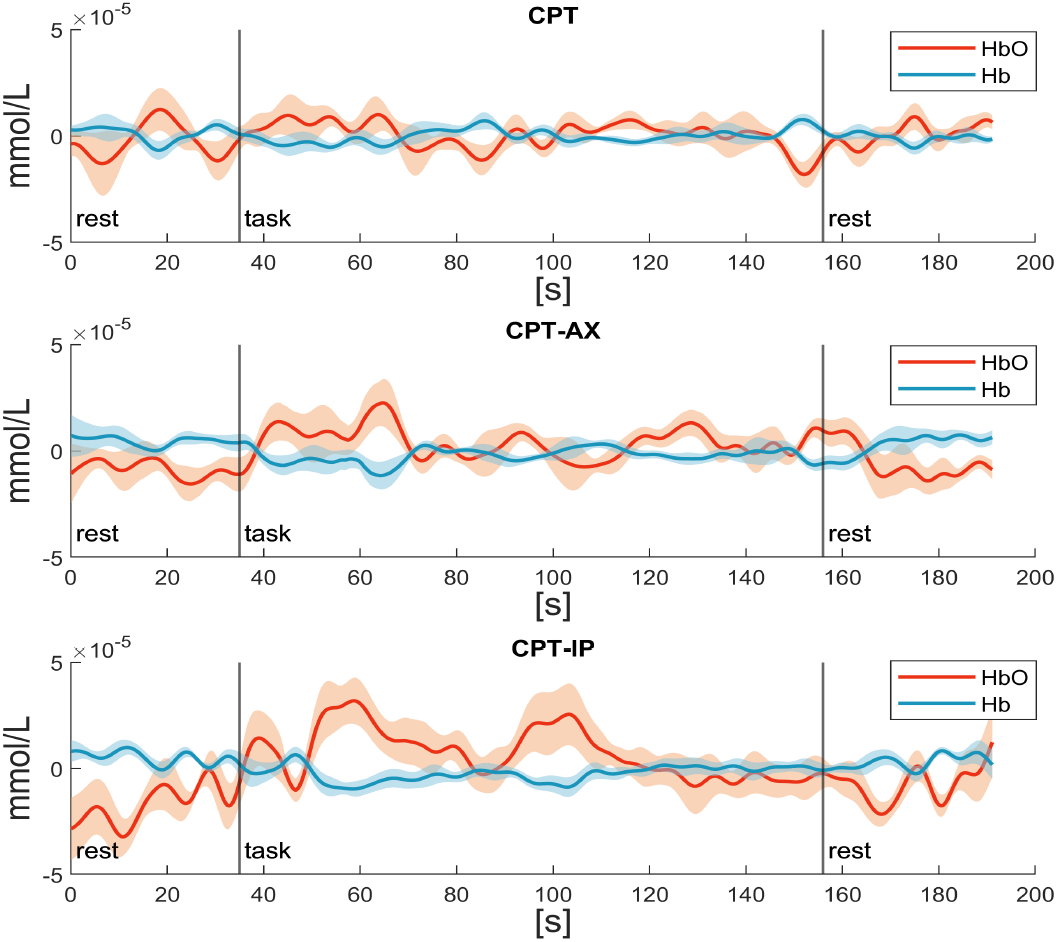
Grand average (solid line) and SEM (shaded area) responses are illustrated for HbO in red and Hb in blue for a) CPT-X CPT-AX c) CPT-IP tests. The two black vertical lines denote the start and end points of the 120s test.

In order to compute heart rate from the fNIRS signal, raw HbO signals of all channels were filtered by an FIR low-pass filter at 1.9 Hz and an FIR High-pass filter at 1 Hz. Hence, unwanted components, including high-frequency motion artifacts and other physiological noises unrelated to the heart rate, were removed.

### HRV derivation

After eliminating unwanted components except the heart rate noise from HbO signals, the algorithm proposed by Hakimi et al.^31^ was used for the HR derivation. This algorithm is a real-time algorithm which was employing a weighted mean of four peak detection algorithms, including AMPD, S1 and S5 functions, and M2. Furthermore, it had an effective correction process to extract a more accurate heart rate from fNIRS signals. By extracting the heart rate from each channel, HRV was calculated. Finally, HRV averaged over all channels for further analysis. Figure 6 indicates the fourth participant’s HRV data during one session. In this figure, red parts correspond to 120 s CPT-IP test blocks, and no-colored parts are related to the rest states, which are at the first and end of the session and between attention test blocks.

**Figure 6.**
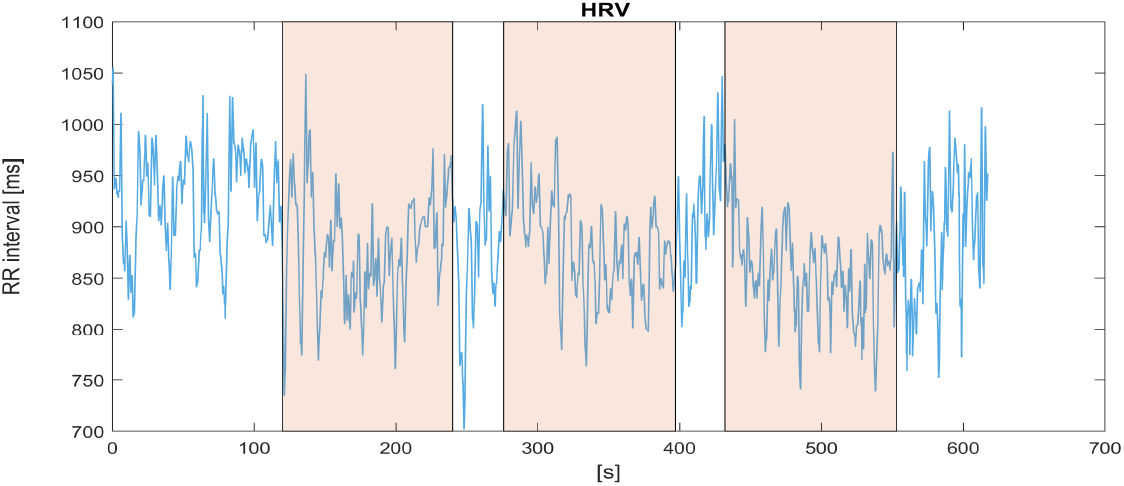
An example of HRV data extracted from the fourth participant’s HbO signals during the CPT-IP test session. The three red parts are related to attention states in which the participant did the attention test, and parts with no color correspond to the rest states.

### Feature extraction

In this study, averaged HbO, Hb, and extracted HRV over all channels were considered for feature extraction, and features were calculated at 15s time windows.

For fNIRS signals, features consisted of three categories: statistical features in the time domain (*m*_*T*_, *Std*_*t*_, *S*_*t*_, *K*_*t*_), statistical features in the frequency domain (*m*_*f*_, *Std*_*f*_)^31^, and morphological features (*NoP, SoP, SS, SP*)^11,42^. Time Statistical features were common features in studies of mental workload and Brain Machine Interface^41,53^. Frequency statistical features also revealed the fluctuation of signals effectively. Table 1 shows fNIRS features and their descriptions.

**Table 1.** Features extracted from fNIRS signals (HbO and Hb) and their descriptions.

| fNIRS features | Description |
| --- | --- |
| Mean ($m_t$) | $\frac{1}{N} \sum_{i=1}^N x_i$ |
| Standard deviation ($Std_t$) | $\sqrt{\frac{1}{N} \sum_{i=1}^N (x_i - m_t)^2}$ |
| Skewness ($S_t$) | $\frac{1}{N * Std_t^3} \sum_{i=1}^N (x_i - m_t)^3$ |
| Kurtosis ($K_t$) | $\frac{1}{N * Std_t^4} \sum_{i=1}^N (x_i - m_t)^4$ |
| Frequency mean ($m_f$) | $\sum_{i=1}^N X'(f_i) * f_i$ |
| Frequency standard deviation ($Std_f$) | $\sqrt{\sum_{i=1}^N (f_i - m_f)^2 * X'(f_i)}$ |
| Number of peaks ($NoP$) | The total number of local maxima of signals in the selected time windows. |
| Sum of peak ($SoP$) | The sum of the maxima values in the selected time window. |
| Signal slope ($SS$) | The slope of the regression line fits the data in the selected time window using the polyfit function in MATLAB. |
| Signal peak ($SP$) | The peak of data in the selected time window estimated by the MATLAB max function. |

In table 1, *x*_*i*_: *i* = 1,…, *N* is a windowed signal with the desired length. The signal *X′*(*f_i_*); *i* = 1,..., *N* is normalized Fourier transform amplitude and *f*_*i*_ is the digitized frequency in the range of [0,*Fs/*2], which *Fs* is the sampling frequency of the signal. *X′*(*f_i_*) is calculated according to Eq. (1).

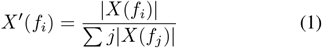

For HRV data, features are divided into time, frequency, and nonlinear domains. Time domain features considered in this study were *MeanNN, StdNN*, and *RMSSD*^54,55^. Frequency domain features selected were categorized into statistical features including *Mean*_*f*_ *NN* and *Std*_*f*_ *NN* Fourier transform of NN intervals and features based on power spectral density of the HRV including *LF, HF*, and *LF/HF*^14,31^. Furthermore, *SD*1, *SD*2, and *SD*1*/SD*2 were used as three nonlinear features obtained from the Poincare plot of HRV, in which each RR interval is plotted against the prior interval. *SD*1 and *SD*2 are the standard deviations of the Poincare plot perpendicular to and along the line-of-identity, respectively^54,55^. HF reflects parasympathetic activity, and *LF* represents both sympathetic and parasympathetic activity but is more related to sympathetic activity. *RMSSD* and *SD*1 correlate with *HF* . Moreover, *SD*2 correlates with *LF*, and *LF/HF* correlates with *SD*1*/SD*2, which assessed balance between parasympathetic and sympathetic activity (autonomic balance)^54,56^. HRV features and their descriptions are shown in table 2.

**Table 2.** Features extracted from HRV data and their descriptions.

| HRV features | Description |
| --- | --- |
| $MeanNN$ | Mean of NN intervals (calculated like the $m_t$ in table 1) |
| $StdNN$ | The standard deviation of NN intervals (calculated like the $Std_t$ in table 1) |
| $RMSSD$ | Root mean square of successive RR interval differences:<br>$\sqrt{\frac{1}{N-1} \sum_{i=1}^{N-1} (RR_{i+1} - RR_i)^2}$ |
| $Mean_fNN$ | Mean of Fourier transform of NN intervals (calculated like the $m_f$ in table 1) |
| $Std_fNN$ | The standard deviation of Fourier transform of NN intervals (calculated like the $Std_f$ in table 1) |
| $LF$ | Power of low-frequency band (0.04–0.15 Hz) |
| $HF$ | Power of high-frequency band (0.15–0.4 Hz) |
| $LF/HF$ | The ratio of the $LF$ to $HF$ power |
| $SD1$ | $\frac{1}{\sqrt{2}} \sqrt{var(x - y)}$<br>$x = RR_1, RR_2, \dots, RR_{(N-1)}$<br>$y = RR_2, RR_3, \dots, RR_N$ |
| $SD2$ | $\sqrt{\frac{2}{N} \sum_{i=1}^N (x_i - \bar{x})^2 - SD1^2}$<br>$x = RR_1, RR_2, \dots, RR_N$ |
| $SD1/SD2$ | The ratio of the $SD1$ to $SD2$ |

After features extraction, the feature values were normalized between -1 and 1 by Eq. (2).

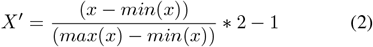

Where x is the original feature value, *x′* is normalized feature value, min(x), and max(x) indicate the smallest and largest feature value.

### Attention classification

After preprocessing and feature extraction, attention classification was done on fNIRS signals (HbO and Hb), HRV data, and both of them simultaneously. The SVM with RBF kernel, LDA, and RF classifiers were employed to distinguish between different sustained attention levels and separate the rest from the attention state. For each classifier, the classification was performed on each participant with 10-fold cross-validation, and the averaged accuracy of participants was reported.

The optimal subset of features was selected by the mutual information (MI) method for reducing classification error. Figure 7 demonstrates the block diagram of the optimal feature selection and classification. For this purpose, 50% of data considered as train and 50% considered as test data, i.e., data of four participants were chosen randomly for training and others were used for testing. Afterward, the feature selection and classification were performed as follows:

**Figure 7.**
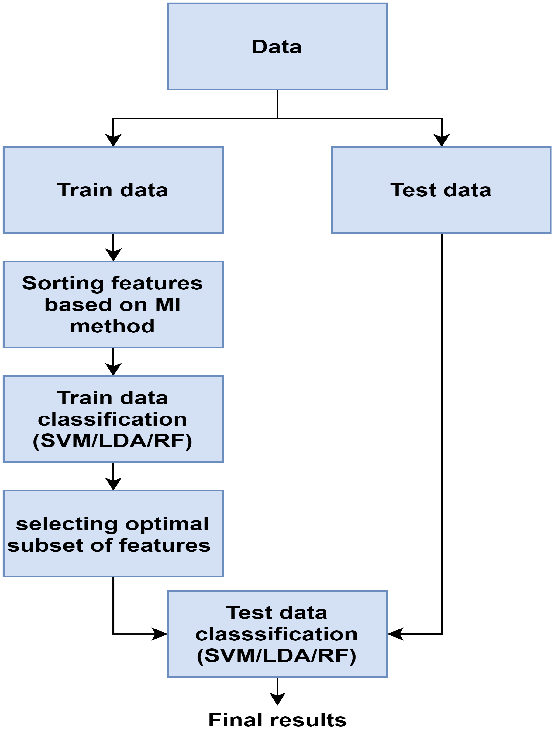
The general flow of selecting the optimal subset of features and classification.

1. For individual participants in the training data set, the mutual information value for each feature was calculated. The features’ MI values averaged over participants. Then features are ranked based on their MI value.
2. Ten features with the highest MI value were selected. The classification accuracy with “n” number of features (n varies between one to 10) was computed.
3. Between 10 selected features, the optimal subset of features was picked where maximum accuracy was reached.
4. Eventually, by determining optimal features, test data were classified.

## Results

### Behavioral data analysis

Figure 8 depicts the average rate of true answers, false alarms, and reaction time during CPT-X, CPT-AX, and CPT-IP. The average rate of true answers decreased from CPT-X to CPT-AX to CPT-IP. In contrast, the average rate of false alarms and the average reaction time increased from CPT-X to CPT-AX to CPT-IP. Repeated measure ANOVA test over these parameters showed that average rate of true answers (F(2,14)=6.36, p-value=0.010), false alarms (F(2,14)=11.58, p-value=0.001), and reaction time (F(2,14)=40.15, p-value*<*0.001) were significantly influenced by the level of task. these results confirmed that participants experienced sustained attention at different levels during CPTs. Pair-wisehow can find comparisons after Bonferroni adjustment showed that there are significant differences in the average rate of true answers (p-value=0.008), rate of false answers (p-value=0.007), and reaction time (p-value*<*0.0001) between CPT-X and CPT-IP (p). Also, there are significant differences in the average rate of false answers (p-value=0.016) and reaction time (p-value*<*0.001) between CPT-AX and CPT-IP. Furthermore, averaged attention scores reported by participants between 1 to 10 were 2.75±2.6,4.87±2.1, and 7.86±1.35 during CPT-X, CPT-AX, and CPT-IP, respectively. This result also indicated that each CPT demanded a different level of sustained attention.

**Figure 8.**
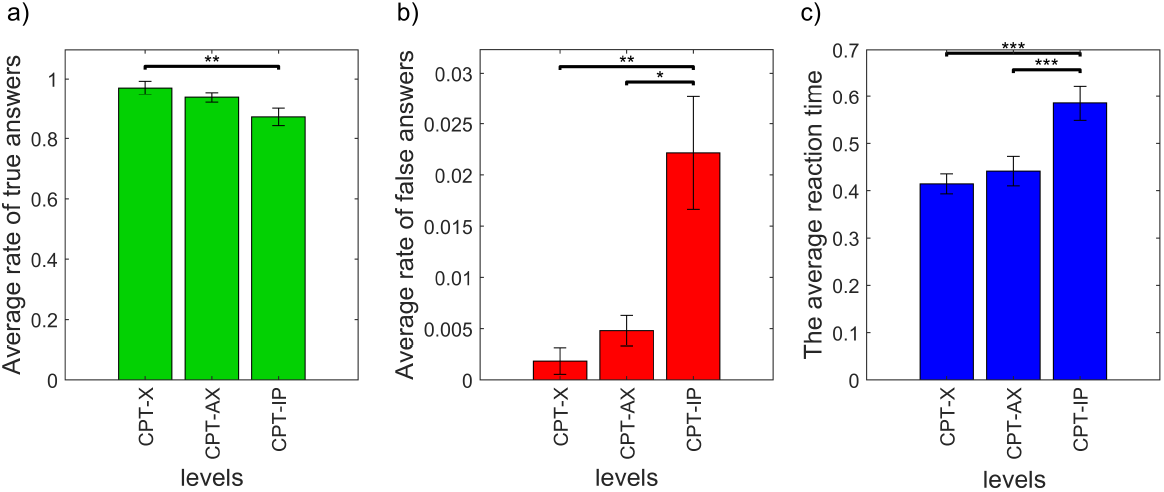
a) Average rate of true answers achieved in different attention levels corresponding to CPT-X, CPT-AX, and CPT-IP b) Average rate of false alarms achieved in different attention levels. c) Average reaction time of true answers achieved in different attention levels. *, **, and *** show significantly difference in which p-value*<*0.05, p-value*<*0.01, and p-v*<*0.001, respectively.

### Result of attention lassification

Table 3 demonstrates classification accuracies by using fNIRS, HRV, and both fNIRS and HRV features with SVM, LDA, and RF classifiers. To analyze and compare fNIRS and HRV data’s classification performance, three-class classification accuracy related to three levels of sustained attention-which were including CPT-X, CPT-AX, and CPT-IP tests- and classification accuracies of all pair of sustained attention levels were obtained for each classifier. Moreover, the optimal subsets of features selected by the MI method are shown in table 3 for each classification condition. The highest three-class classification accuracy was 80.9±1.5, 56.2±1.0, 84.3±0.7 obtained by LDA classifier using fNIRS, HRV, and both fNIRS and HRV features, respectively. Pairwise classification accuracies in table 3 indicate (CPT-X and CPT-IP) were more discriminative than (CPT-X and CPT-AX) and (CPT-AX and CPT-IP). Furthermore, fNIRS features could classify attention levels more accurately than HRV features, and using both of them simultaneously increased classification accuracies in all classifiers. According to selected features columns in table 3, among fNIRS features, all classifiers used (*SP*)_*HbO*_, (*M*_*t*_)_*HbO*_, (*Std*_*f*_)_*HbO*_, (*Std*_*f*_)_*Hb*_ and between HRV features, all classifiers used *LF* power. So, these features were more discriminative features than others.

**Table 3.** Classification accuracies (%) and the optimal subset of features for classifying sustained attention into three and two states with SVM, LDA, and RF classifiers by employing fNIRS features, HRV features, and both of them.

| Classification accuracies using fNIRS features |  |  |  |  |
| --- | --- | --- | --- | --- |
| Task | SVM | LDA | RF | Selected Features |
| 3 Class Classification | $71.1 \pm 2.5$ | **$80.9 \pm 1.5$** | $68.1 \pm 3.2$ | SVM/LDA/RF: $(SP)_{HbO}$, $(M_t)_{HbO}$, $(Std_f)_{HbO}$, $(Std_t)_{HbO}$, $(Std_f)_{Hb}$ |
| CPT-X vs. CPT-AX | $84.9 \pm 1.9$ | **$94.1 \pm 1.1$** | $81.9 \pm 3.5$ | SVM/LDA/RF: $(SP)_{HbO}$, $(M_t)_{HbO}$, $(Std_f)_{HbO}$, $(Std_t)_{HbO}$, $(Std_f)_{Hb}$ |
| CPT-AX vs. CPT-IP | $90.5 \pm 0.9$ | **$93.2 \pm 1.8$** | $87.0 \pm 2.4$ | SVM/LDA/RF: $(SP)_{HbO}$, $(M_t)_{HbO}$, $(Std_f)_{HbO}$, $(Std_f)_{Hb}$ |
| CPT-X vs. CPT-IP | $92.1 \pm 2.2$ | **$93.7 \pm 1.8$** | $85.9 \pm 0.8$ | SVM/LDA/RF: $(SP)_{HbO}$, $(M_t)_{HbO}$, $(Std_f)_{HbO}$, $(Std_f)_{Hb}$ |
| Classification accuracies using HRV features |  |  |  |  |
| Task | SVM | LDA | RF | Selected Features |
| 3 Class Classification | $55.5 \pm 2.3$ | **$56.2 \pm 1.0$** | $55.5 \pm 3.5$ | SVM: $LF$<br>LDA: $LF$, $SD1$, $LF/HF$, $RMSSD$<br>RF: $LF$, $SD1$, $LF/HF$, $RMSSD$ |
| CPT-X vs. CPT-AX | **$71.1 \pm 3.6$** | $69.9 \pm 2.0$ | $69.6 \pm 2.2$ | SVM: $LF$<br>LDA: $LF$, $SD1$, $LF/HF$<br>RF: $LF$, $SD1$, $LF/HF$, $RMSSD$ |
| CPT-AX vs. CPT-IP | $63.7 \pm 2.3$ | $67.7 \pm 1.2$ | **$70.8 \pm 1.9$** | SVM: $LF$<br>LDA: $LF$, $SD1$, $LF/HF$, $RMSSD$<br>RF: $LF$, $SD1$, $LF/HF$ |
| CPT-X vs. CPT-IP | **$71.1 \pm 3.3$** | $70.3 \pm 2.3$ | $67.7 \pm 1.8$ | SVM: $LF$<br>LDA: $LF$, $SD1$, $RMSSD$, $Std_fNN$, $StdNN$<br>RF: $LF$, $SD1$, $RMSSD$, $Std_fNN$ |
| Classification accuracies using both fNIRS and HRV features |  |  |  |  |
| Task | SVM | LDA | RF | Selected Features |
| 3 Class Classification | $71.7 \pm 2.1$ | **$84.3 \pm 0.7$** | $78.1 \pm 1.9$ | SVM/LDA/RF: $(SP)_{HbO}$, $(M_t)_{HbO}$, $(Std_f)_{HbO}$, $LF$, $(Std_t)_{HbO}$, $(Std_f)_{Hb}$ |
| CPT-X vs. CPT-AX | $86.5 \pm 2.4$ | **$94.5 \pm 1.5$** | $86.3 \pm 2.3$ | SVM/LDA/RF: $(SP)_{HbO}$, $(M_t)_{HbO}$, $(Std_f)_{HbO}$, $LF$, $(Std_t)_{HbO}$, $(Std_f)_{Hb}$ |
| CPT-AX vs. CPT-IP | $94.9 \pm 1.7$ | **$95.2 \pm 1.5$** | $95.0 \pm 1.1$ | SVM/LDA/RF: $(SP)_{HbO}$, $(M_t)_{HbO}$, $(Std_f)_{HbO}$, $LF$, $(Std_t)_{HbO}$, $(Std_f)_{Hb}$ |
| CPT-X vs. CPT-IP | $82.1 \pm 1.3$ | **$95.6 \pm 1.2$** | $90.5 \pm 1.6$ | SVM/LDA/RF: $(SP)_{HbO}$, $(M_t)_{HbO}$, $(Std_f)_{HbO}$, $LF$, $(Std_f)_{Hb}$ |

Table 4 shows the rest against attention classification accuracy by fNIRS, HRV, and both fNIRS and HRV features. Different attention levels were considered one class to differentiate between the rest and attention state. The RF classifier had the best performance in classifying rest and attention state. Also, using both fNIRS and HRV features increased RF classification accuracy to 99.5±0.2. In contrast, combining fNIRS and HRV features did not improve SVM classification accuracy in this situation. For classifying rest and attention, (*SP*)_*HbO*_ and (*NoP*)_*Hb*_ features from fNIRS signals and *LF* power from HRV data had the highest MI value between features.

**Table 4.** Classification accuracies (%) and the optimal subset of features for classifying sustained attention into the attention and rest state with SVM, LDA, and RF classifiers by employing fNIRS features, HRV features, and both of them.

| Attention vs. Rest classification accuracies |  |  |  |  |  |
| --- | --- | --- | --- | --- | --- |
| Type of features | SVM | LDA | RF | Selected Features |  |
| fNIRS features | 95.1 ± 0.8 | 72.5 ± 0.1 | **98.9 ± 0.3** | SVM: $(SP)_{HbO}$, $(NoP)_{Hb}$, $(NoP)_{HbO}$, $(Std_f)_{HbO}$<br>LDA: $(SP)_{HbO}$, $(NoP)_{Hb}$<br>RF: $(SP)_{HbO}$, $(NoP)_{Hb}$, $(NoP)_{HbO}$ | |
| HRV features | 83.0 ± 1.1 | 77.4 ± 1.1 | **95.6 ± 1.2** | SVM: $LF$<br>LDA: $LF$, $SD1/SD2$, $HF$, $MeanNN$, $StdNN$<br>RF: $LF$ | |
| fNIRS and HRV features | 81.7 ± 0.9 | 79.7 ± 1.2 | **99.5 ± 0.2** | SVM: $(SP)_{HbO}$, $(NoP)_{Hb}$, $LF$, $(NoP)_{HbO}$<br>LDA: $(SP)_{HbO}$, $(NoP)_{Hb}$, $LF$<br>RF: $(SP)_{HbO}$, $(NoP)_{Hb}$, $LF$, $(NoP)_{HbO}$ | |

## Discussion

The aim of this study was to investigate the feasibility of fNIRS signals and HRV data derived from fNIRS signals in classifying sustained attention states in three levels and discriminate the rest from sustained attention state. The fNIRS device measured eight participants’ hemodynamic response during rest and three different versions of the CPT (CPT-X, CPT-AX, and CPT-IP). Then, the HRV data was derived by the weighted mean algorithm from the HbO signal. After applying signal processing algorithms, specified features were extracted from fNIRS and HRV data. The feature selection algorithm based on MI determined the optimal subset of features for each classifier. SVM, LDA, and RF classifiers were used to classify sustained attention by fNIRS, HRV, and both fNIRS and HRV features. Due to individual variations in the hemodynamic response and heart rate, classifications were done on each participant and averaged accuracy over participants was considered. The results are then explained.

Behavioral results during CPTs indicate the significant decrement in the rate of true answers and the significant increment in the rate of false alarms and the reaction time from CPT to CPT-AX to CPT-IP. These results show that CPTs can induce different sustained attention levels, which corresponds to previous studies^6,57^. Furthermore, subjective scales also show different attention loads in each CPT.

fNIRS devices measure the hemodynamic response of the brain contaminated with physiological noises like the heart rate. Most of the past research used fNIRS signals for studying attention, but none of them used HRV data extracted from fNIRS signals^4,12,13,58^. Therefore, We used fNIRS and HRV extracted from fNIRS signals for assessing both heart and brain’s behavior during sustained attention. The best three-class sustained attention classification accuracy with fNIRS features was 80.9% by LDA classifier that is 23.05% higher than the previous study reaching for three-class sustained attention classification^16^. This achievement shows the efficiency of MI based feature selection and preprocessing algorithms. The HRV features reached 56.2% accuracy for three-class classification by LDA classifier, which confirming the brain and heart interactions and the ability of HRV for evaluating sustained attention. Consequently, by fusing both fNIRS and HRV features, three-class accuracy becomes 84.3% with LDA classifier, higher than fNIRS and HRV alone. Binary classification accuracy between each level of sustained attention corresponding to CPTs shows that CPT-X and CPT-AX were less discriminative than other pairs. In summary, fNIRS features were more capable of classifying different attention levels than HRV features. Also, combining fNIRS and HRV features increased the classification accuracy of all classifiers.

The classification result between rest and attention states represent HRV features can differentiate these states as good as fNIRS features. The best classification accuracies achieved through the RF classifier are 98.9% and 95.6% by fNIRS and HRV features. For classification between rest and attention, only the RF classifier had a better performance by fusing features, In contrast to classification between different sustained attention levels.

Between different levels of sustained attention, the MI based feature selection algorithm indicates that the signal peak and mean value of the HbO signal have higher MI values than other features that are in accordance with other studies finding these two features more discriminative^42,43^.

For the HRV data, *LF* and *SD*1*/SD*2 have the higher MI value in which *LF* corresponds to the sympathetic nervous system, and *SD*1*/SD*2 shows the autonomic balance. Between these two features, *LF* is frequently used in past research for evaluating cognitive functions^55,59,60^.

One limitation of the current work is the small number of participants. Referring to past studies, eight participants are enough for evaluating cognitive tasks^42,51^. However, a larger sample size might be helpful for results validation. Furthermore, by more participants, other validation algorithms such as the leave-one-out can be applied to measure the classification performance independent of the subject^61^. Another limitation of this study is considering only females with an average age of 23.8± Classification accuracies using using both fNIRS and HRV features 0.97 for data acquisition because gender and age influence HRV features. Hence, it is necessary to consider both male and female subjects of different ages for future work to investigate the effect of gender and age on classification accuracy and features^54,62^. Finally, the best features chosen for evaluating sustained attention by the fNIRS and HRV data derived from the fNIRS brain signals can be used in biofeedback systems considering both brain and heart response to improve attention for people with attention disorders.

## Conclusion

in this study, we examined the capability of fNIRS and HRV data extracted from fNIRS signals to classify sustained attention levels induced by three different versions of CPTs. Also, the study investigated the potential of these signals to classify the rest from the attention state. Our results showed fNIRS signals were more capable than HRV data to classify three or two sustained attention levels. However, fNIRS signals and HRV data had the same potential to differentiate rest from attention states. Moreover, combining both fNIRS and HRV features increased classification accuracies. This result suggests that HRV extracted from fNIRS signals can be used with the hemodynamic response to increased multiclass classification performance for discriminating different cognitive states such as sustained attention levels. Hence, one fNIRS device provides more accurate biofeedback or Brain-Computer Interface (BCI) systems without adding additional modalities.

## Acknowledgments

The authors would like to appreciate all students who participated as subjects in this experiment.

## Declaration of competing interest

The authors declare that they have no known competing financial interests or personal relationships that could have influenced the work reported in this paper.

## Notes

### Competing Interest Statement

The authors have declared no competing interest.

